# Ancient nervous system architecture in a living ctenophore

**DOI:** 10.64898/2026.05.20.726345

**Authors:** Anna Ferraioli, Paula Miramón-Puértolas, Paula Emilia Altenkirch, Alexandre Jan, Jeffrey Colgren, Jakob Vinther, Pawel Burkhardt

## Abstract

The evolutionary origin of nervous systems in animals remains elusive and is largely hidden from the fossil record. Ctenophores, one of the earliest-branching animals possessing neurons, are instrumental to our understanding of nervous system origin, and a few rare ctenophore fossils preserve traces of nervous tissue as carbonaceous remains. Cambrian ctenophores appear to exhibit a more diverse neuroanatomy than that of modern species, suggesting secondary loss in extant ctenophores. However, much remains unknown about the origin and ontogeny giving rise to the structural organization of modern ctenophore nervous systems. Here, by investigating the neural anatomy of the ctenophore *Mnemiopsis leidyi* during development, we identified a ladder-like nerve net (LNN) beneath the comb rows that converges into condensed neurites and connects to the aboral organ. Examination of carbon-rich areas of *Ctenorhabdotus capulus*, an extinct ctenophore from the Burgess Shale, reveals a pattern similar to that of *M. leidyi*, consistent with a shared neural organization. Furthermore, *M. leidyi* exhibits a condensed comb nerve, resembling the longitudinal nerve preserved in the Cambrian ctenophore *Fasciculus vesanus* and the giant axon of extant *Euplokamis dunlapae*. Our study reveals conserved evolutionary constraints shaping nervous system architectures linked to locomotory organs and indicates that the different modes of nervous system organization observed in Cambrian ctenophores are variably retained in modern species.

## Introduction

Nervous systems exhibit remarkable diversity across animals, ranging from diffuse nerve nets to highly centralized brains. Understanding how these different architectures evolved remains a central challenge in evolutionary biology (1–3). Comparative studies suggest that nervous system organization is shaped not only by lineage-specific histories but also by shared functional and developmental constraints. Identifying neural architectures in early-branching animal lineages can therefore provide key insights into the principles governing the emergence and diversification of nervous systems.

Various attempts at reconstructing the early-metazoan phylogenetic history have placed ctenophores as the earliest-branching animals (4–8). Although still under debate (9,10), this finding would support the scenario of multiple origins of neurons (4,8,11). A less well-supported hypothesis recovered in some molecular studies (12– 14) and supported by morphology and the fossil record (15), places ctenophores as a sister taxon to Cnidaria, implying a single origin of nervous systems in which ctenophores underwent major secondary transformations at the genetic level. Regardless of their placement, ctenophores remain essential for understanding the evolution of neurons and nervous systems (11).

Ctenophores experienced a major species diversity bottleneck (16,17) hundreds of millions of years after their divergence, suggesting that extant species represent only a small fraction of their original diversity, which is a critical consideration when reconstructing their origin and the history of ancestral traits. However, several stem ctenophores from the Cambrian have been found. Exceptionally preserved fossils have revealed traces of nervous systems (18–24), found as carbon-rich remains from a diversity of animals (15,19,20,25,26). These discoveries provide rare access to neuroanatomy during the Cambrian Explosion and offer a critical framework for reconstructing the evolutionary history of early-branching lineages (15,25). Notably, neural elements are inferred to be preserved in the ctenophores from the early and middle Cambrian (15,25), providing key evidence for investigating the deep origins of their nervous systems.

Soft body parts, including comb row cushion plates, nervous systems, elements of the aboral organ, and sclerotized organic skeletons lost in living ctenophores, are documented in Burgess Shale-type Lagerstätten, facilitating the analysis of evolutionary relationships, anatomy, and body plan transformations (15,25,27,28). Cambrian ctenophores exhibit diverse symmetries in which sessile stem lineages exhibit a triradial configuration with six to 18 tentacles (15), while scleroctenophores exhibit eight paired comb rows (28), and the slightly younger taxa known from the Burgess shale exhibit 24 (eight trifurcating comb rows) to more than 80 comb rows (27). Cambrian ctenophores, with apparent pelagic modes of life, all exhibit a protruding aboral organ, often encapsulated, within a skeletal cone in scleroctenophores, and flaring oral lobes or an ‘oral skirt’ (25). A possible ‘circumoral nerve ring’ around the oral opening is a feature reported in the Cambrian species *Thalassostaphylos elegans* and *Ctenorhabdotus campanelliformis* (25). A longitudinal nerve beneath the comb rows is present in *C. campanelliformis* and *Fasciculus vesanus*, whereas the scleroctenophore *Galeactena hemispherica* features a longitudinal nerve extending between the paired comb rows underneath a supportive skeletal rod (15,25). The elaborate and diverse neuroanatomical conditions in Cambrian fossils, in contrast to those in extant species, suggest that aspects of that complexity were lost before the divergence of extant ctenophores (25).

In modern species, the nervous system typically consists of a polygonal subepithelial nerve net (SNN) (29–33) including distinctive neural elements beyond the SNN (Fig. S1). *Euplokamis dunlapae*, the sister group to all other ctenophores (34), possesses a prominent comb nerve or axon that runs longitudinally beneath the comb rows, linking adjacent comb plates - a feature unique to this lineage (33,35). Tentacular nerves extend from the aboral organ toward the tentacles in *E. dunlapae* and *Pleurobrachia* sp. (31,33,36) (Fig. S1). Benthic species such as *Vellicula multiformis* also show the polygonal nerve net (29,33,37). Finally, a “giant lip nerve” encircles the specialized oral structure unique to Beroids (32,38) (Fig. S1). Recent work has revealed hitherto unknown molecular components to the ctenophore nervous system, including a host of unique neuropeptides, which provide novel markers for exploring the development and localization of different components of the ctenophore nervous system (4,39–41).

Here, we map the nervous system architecture of the ctenophore *Mnemiopsis leidyi* from juveniles to adults using expression patterns of neuropeptide precursors. During development, the polygonal nerve net condenses and becomes highly organized within the locomotory comb row region. Spatial segregation of gene expression and condensation of neurites indicate the emergence of distinct neural circuits in those organs. Remarkably, the resulting neuroanatomical patterns mirror the carbonaceous imprints of the Cambrian fossil *Ctenorhabdotus capulus*, suggesting that key elements of modern ctenophore neural architecture have deep evolutionary origins.

## Results

### Ladder-like nerve net (LNN) architecture and spatial segregation of neuropeptide expression

We examined nerve net architecture during development by mapping the expression of the *Mnemiopsis*-specific neuropeptide precursors ML199816a and ML02212a (39) using hybridisation chain reaction (HCR; Fig. 1). Both genes are broadly expressed in both neural cell bodies and neurites, enabling visualization of the ctenophore nerve net architecture (Fig. 1, Movie S1). During the first week of development, the polygonal SNN below the comb rows of post-hatching animals (29) converts into a ladder-like nerve net (LNN) (Fig. 1A, B, Fig S2, Movie S1). Neurites and cell bodies are located between polster cell clusters (Fig. 1A, B, S2A) and co-express the two neuropeptide precursors (Fig. 1A, B, C). Longitudinal neurites run along the eight comb rows (Fig. 1A, B), while transverse neurites connect these at the oral and aboral margins of the comb plates (Fig. 1A, B, S2A). One to two cell bodies expressing both neuropeptide precursors are present on the transverse neurites, whereas almost no cell bodies are observed across the longitudinal ones (Fig. 1A, B). Cells in the aboral organ and tentacle bulbs are enriched with the expression of *ML02212a* solely (Fig. 1A, B). In contrast, cells forming a nerve tract in the mid-part of the tentacle bulb are enriched with the expression of *ML199816a* (Fig. 1A). This nerve tract is defined as the tentacle nerve. Neurites originating within the aboral organ connect to the LNN of the eight comb rows (Fig. 1A, Movie S1), whereas the LNN remains associated with the broader polygonal nerve net (Fig. 1C).

**Figure 1.**
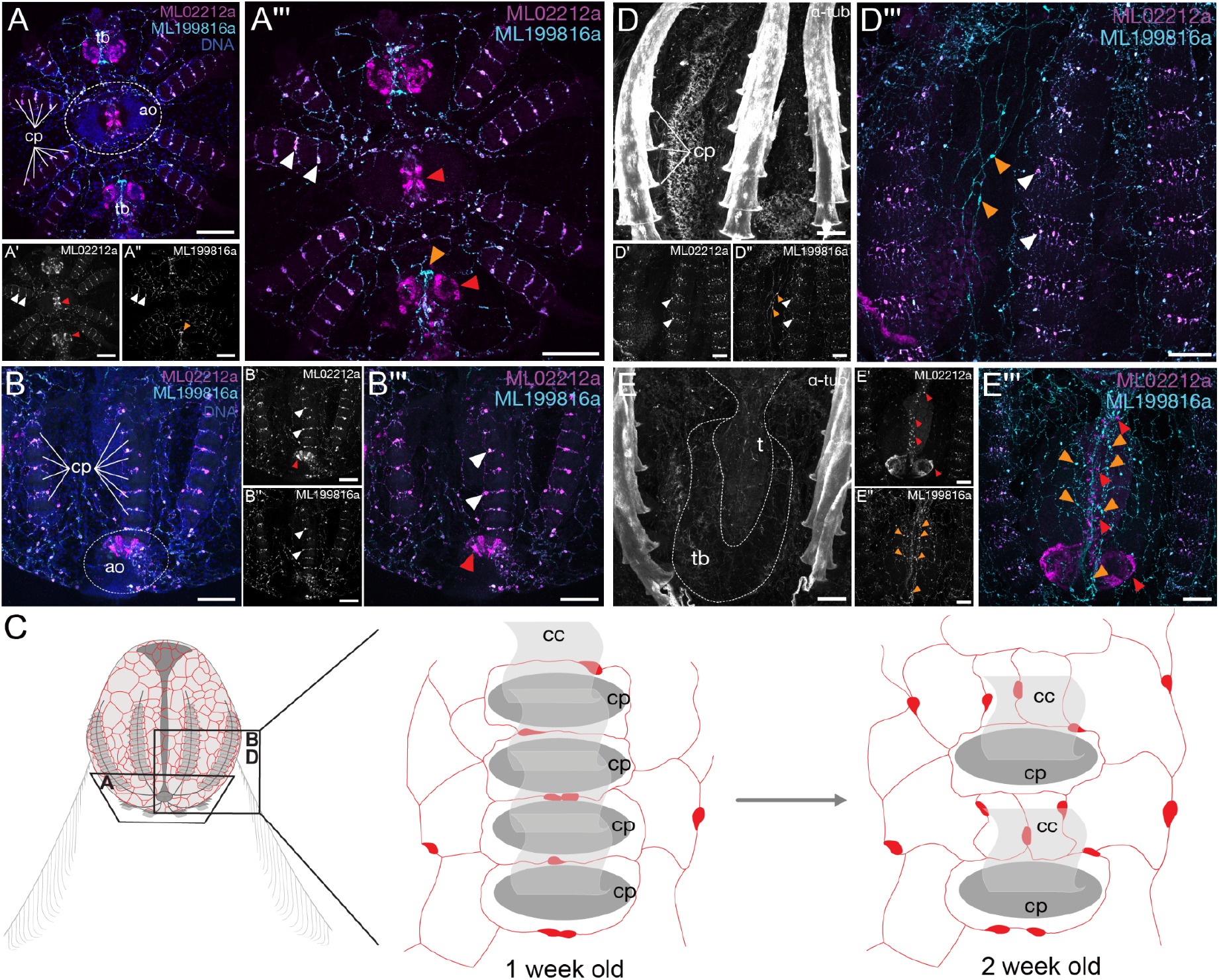
Neuropeptide expression patterns reveal ladder-like (LNN) architecture and spatial segregation of gene expression in the nerve net. HCR showing the expression of ML02212a and ML199816a in one-week-old (A, B) and two-week-old cydippids (D, E) of M. leidyi. (A) Aboral view and (B) lateral view. The expression of ML02212a (magenta) and ML199816a (cyan) highlights a ladder-like nerve net (LNN) architecture formed between comb plates (cp). DNA is stained with Hoechst, and nuclei are shown in blue. (A’, B’) Expression of ML02212a. (A’’, B’’) Expression of ML199816a. (A’’’, B’’’) Merged expression of ML02212a and ML199816a. Lateral view of comb rows (D) and tentacle (E), cilia are stained with anti-a-tubulin antibody (grey). (D’, E’) Expression of ML02212a. (D’’, E’’) Expression of ML199816a. (D’’’, E’’’) Merged expression of ML02212a (magenta) and ML199816a (cyan). White arrowheads indicate co-localization, red arrowheads indicate enriched expression of ML02212a, and orange arrowheads indicate enriched expression of ML199816. (C) Schematic of a cydippid (left panel) and the developmental progression of the nerve net architecture aligning with the comb plates (cp) after one (mid panel) and two weeks (right panel). Comb cilia (cc), Aboral organ (ao), Tentacle bulb (tb). Scalebars, 50 µm.

Following two weeks of development, the comb plates become more widely spaced (Fig. 1D, S2B), with several neural cell bodies developing between them (Fig. 1C, D, S2B). At this stage, the transverse component of the LNN consists of multiple cells co-expressing both neuropeptides and interconnected by neurites (Fig. 1D). Longitudinal neurites remain present and appear to be more prominent in between comb rows (Fig. 1D). These express solely *ML199816a* (Fig. 1D). In the tentacle bulb, the tentacle nerve shows strong neuropeptide segregation at this stage (Fig. 1E). *ML02212a* is primarily expressed in cell bodies along the tentacle midline and in two proximal circular bundles within the tentacle bulb (Fig. 1E). *ML199816a*-positive cells form a contiguous tract along the midline of the tentacle bulb and tentacle, with their neurites forming a typical polygonal nerve net (Fig. 1E, 1E).

### Condensed nerve net neurons highlight areas of connectivity

In larger cydippids (∼0.5 mm), the nerve net expands into a larger and more elaborate network by increasing neural cell bodies and neuropeptide segregation (Fig. 3). Expression of the neuropeptide precursors ML02212a and ML199816a marks neurites that converge into condensed structures within defined domains, here termed the ‘ciliated groove nerve’ and the ‘comb nerve’ (Fig. 2A-C, F). The ciliated groove nerve forms beneath the ciliated groove cells at the aboral pole and arises from the convergence of neurites projecting from the aboral organ. (Fig. 2A). These condensed structures predominantly express ML199816a (Fig. 2A), whereas ML02212a is expressed in specific cells of the aboral organ and in a subset of neural cell bodies within the aboral polygonal nerve net (Fig. 2A). The polygonal nerve net surrounding the aboral organ is enriched in ML199816a expression (Fig. 2A). The ciliated groove nerve are continuous with the first comb plate, maintaining strong enrichment of *ML199816a* (Fig. 2B). Proximal to the comb plate, additional expression of *ML02212a* is detected (Fig. 2B) *ML02212a* is also prominently expressed in neural cell bodies surrounding the comb plates (Fig. 2B).

**Figure 2.**
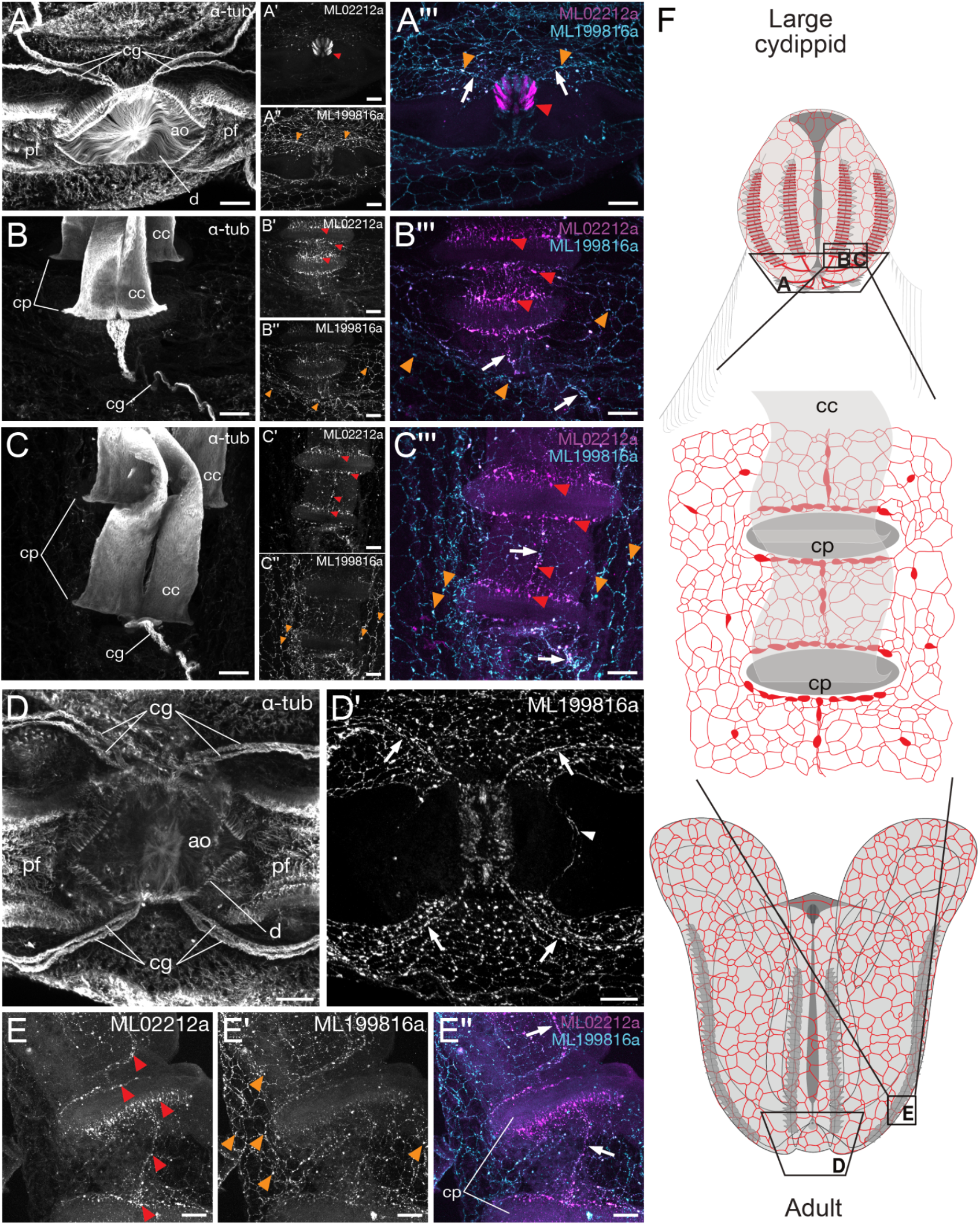
The LNN condenses in larger cydippids and adults. HCR showing the expression of ML02212a and ML199816a combined with tubulin staining in larger cydippids (A, B, C) and adults (D, E) of M. leidyi. (A) Aboral view - aboral organ and (B, C) lateral view of proximal comb plates, including the ciliated groove nerve and the comb nerve. (A, B, C) Cilia are stained with anti-a-tubulin antibody (grey). (A’, B’, C’) expression of ML02212a. (A’’, B’’, C’’) expression of ML199816a. (A’’’, B’’’, C’’’) Merged expression of ML02212a (magenta) and ML199816a (cyan). (D) aboral view of the aboral organ; cilia are stained with anti-a-tubulin antibody (grey). (D’) expression of ML199816a (grey). (E, E’, E’’) Expression of ML02212a (E, grey), ML199816a (E’, grey), and merged expression (E’’, ML02212a in magenta, ML199816a in cyan) in the combs. White arrows indicate condensed neurons forming the ciliated groove nerve in (B’’’) and (D’) and the comb nerve in (C’’’) and in (E’’). White arrowhead in D’ indicates neurites forming a ring-like structure below the aboral organ. Red arrowheads indicate the enriched expression of *ML02212a*, and orange arrowheads indicate the enriched expression of *ML199816a*. (F) Schematic of a large cydippid (top panel), the nerve net architecture around the comb plates (cp) (mid panel), and adult (bottom panel). ao: aboral organ; d: dome; cg: ciliated grooves; pf: polar fields. cp: comb plates; cc: comb cilia. Scalebars, 50 µm.

**Figure 3.**
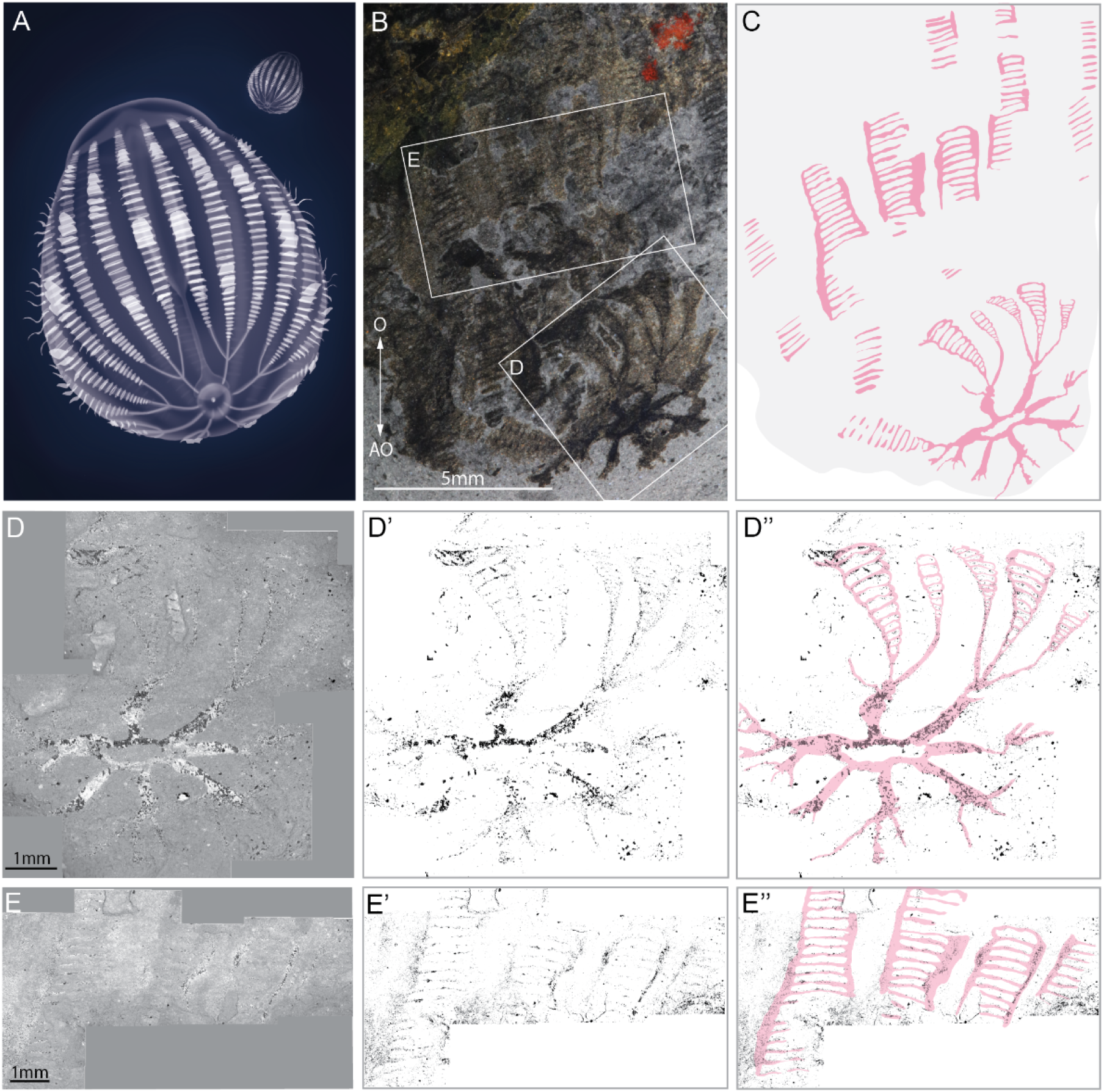
*Ctenorhabdotus capulus*, ROMIP 51439. (A) Schematic 3D reconstruction of *C. capulus*. (B) Complete specimen in lateral view photographed using cross-polarized light (B). (D, E) Scanning electron microscopy backscatter of the aboral organ region (D) and four combs in the mid-body (E). (D’, E’) Scanning electron microscopy images processed to show only the dark elements, enriched in carbon. (C, D’’, E’’) Interpretative drawing (pink) of whole specimen (C), aboral region (D’’) and comb rows (E’’), based on the patterns shown by polarized light and scanning electron microscopy. Original fossil images in B, D, and E are courtesy of J. B. Caron, Royal Ontario Museum.

At this developmental stage, the LNN undergoes spatial rearrangements. The comb plates are more widely spaced and interconnected by ciliated cells forming a tract and named ‘interplate ciliated grooves’ (42,43) (Fig. 2F). The comb nerve emerges as a condensed structure located beneath the interplate ciliated grooves, predominantly expressing *ML02212a* (Fig. 2C, F). In adults of *M. leidyi*, neurites form a ring-like structure beneath the dome at the aboral pole, which is enriched in *ML199816a* expression (Fig. 2D). Condensed ciliated groove nerves (Fig. 2D) and comb nerves (Fig. 2E) are more prominent than in earlier stages, although they share a similar architecture (Fig. 2F, S2C). Despite these important rearrangements, the polygonal nerve net architecture is preserved in later stages of *M. leidyi*, largely defined by the expression of *ML199816a* (Fig. 2B, C, D, E).

### Ladder-like architecture in a ctenophore fossil

Building on the previous discovery of fossilized nervous tissue in ctenophores (15,25,27) and newly described LNN in *M. leidyi*, we explored the ctenophore fossil record to assess whether this neural architecture could be evolutionarily conserved. *Ctenorhabdotus capulus*, a ctenophore from the middle Cambrian Burgess Shale (27), exhibits a wide mouth opening and a body covered by 24 densely spaced comb rows and a protruding aboral organ (Fig. 3A, B, S3) (27). ROMIP 51439 (Fig. 3B, S3) was folded prior to compaction, exposing the surface of the aboral region particularly well (Fig. S3A). The specimen preserves strands of carbonaceous tissue defining the comb rows, which narrow aborally into thin strands (Fig. S3B, C). These strands merge into eight sets, extending to a circular structure that would have been situated underneath the aboral organ (Fig. 3A, B, S3B). Previously, the carbonaceous impressions defining the comb rows were interpreted as the remains of polster cells (15), commonly inferred in other ctenophore fossils. Putative polster cells or cushion plates preserve as discrete, oblong carbonaceous impressions (15,25,28). This is not the case in ROMIP 51439. We comprehensively analyzed the carbon-rich regions in *C. capulus* (Fig. 3A), including dSLR photos of the specimen submerged in water and illuminated with crossed polarized light (Fig. 3B) and details obtained with back-scattered electron imaging (Fig. 3D and E), revealing the ring-like structure underneath the aboral pole from which the eight segments extend toward the comb plates (Fig. 3B, C, S3D, E, F). Within the comb regions, the carbon-enriched areas form a ladder-like network of transverse and longitudinal strands, closely resembling the nerve-net architecture described in *M. leidyi* (see Discussion). Based on the trajectory and topology of the carbonaceous remains preserved in ROMIP 51439, we interpret this tissue as a preserved nervous system.

### Fossils inform on the evolution of ctenophore nervous system architecture

Our revised assessment of the trajectory and organ system identity of the carbonaceous remains associated with the combs of *C. capulus* combined with previous observations (Fig. 4A) (25,27), prompted us to interpret this as a well-preserved nervous system. Comparison of the ladder-like architecture in the comb areas of *C. capulus* and the LNN of *M. leidyi* exhibits strong structural similarities (Fig. 4A, 4A’, insets). The ring-like units of the ladder-like putative nerve structure are tightly interconnected in *C. capulus*, as there is a lack of space between the comb plates (Fig. 4A). The LNN of *M. leidyi* is similarly arranged at early cydippid stages (Fig. 1A, 1B). Later in development, the formation of interplate space and interplate ciliated grooves occurs in parallel with a rearrangement of the LNN and the convergence of neurites into a condensed comb nerve (Fig. 2, 4A’). In the fossils of *Fasciculus vesanus, Ctenorhabdotus campanelliformis*, and the scleroctenophore *Galeactena hemispherica*, a comb nerve showing a similar architecture (i.e., with a continuous trajectory below the overlying comb plates) to that of *E. dunlapae* is retained (Fig. S1, 4B) (15,25,27). These are considered homologous to the *E. dunlapae* giant axon (25). In parallel, condensed nerves beneath the ciliated groove are common to *M. leidyi, E. dunlapae* (33), and to *C. capulus*, in which carbonaceous traces connect the aboral organ to the first comb plates (Fig. 4A).

**Figure 4.**
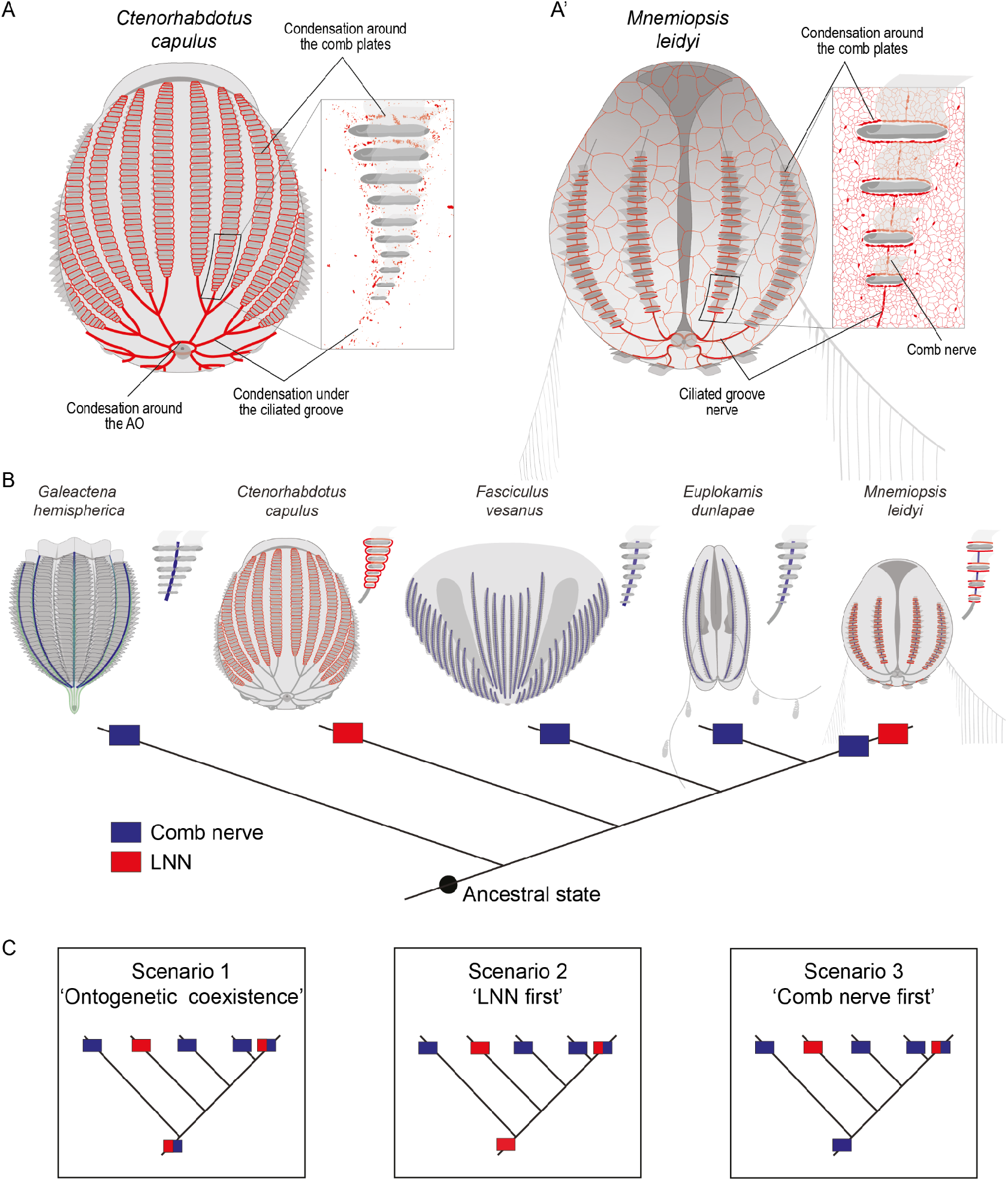
Comparison of fossil and extant ctenophores reveals shared neural features and neural complexity. Diagram of *Ctenorhabdotus capulus* (A) and *Mnemiopsis leidyi (A’)* showing the neural architecture (red) and highlighting the carbonaceous traces and the LNN (insets, red) around the comb plates and underneath the ciliated grooves. (B) Simplified character-based phylogenetic tree including interpretative drawings of fossil species and extant ctenophores with neural features highlighted and insets showing the neural architecture corresponding to the comb rows. The occurrence of carbonaceous ladder-like traces and LNN characters is represented with red squares. The occurrence of the comb nerves is represented with blue squares. The black dot at the base of the tree indicates the open question regarding the ancestral state of the neural architecture. (C) Proposed evolutionary scenarios with hypotheses on the ancestral neural architecture. Scenario 1 depicts the coexistence of the two morphological features and their emergence during ontogeny. Scenario 2 depicts the comb nerve as the ancestral state, and Scenario 3 places the LNN as the ancestral architecture

To determine patterns of nervous system evolution in early and crown group ctenophores, we mapped neural states across a previously published morphologically based phylogeny with *E. dunlapae* fixed as the sister group to all other crown ctenophores (25), and focusing on *G. hemispherica, C. capulus, F. vesanus, E. dunlapae, and M. leidyi* from which aspects of their comb nervous system is known (Fig. 4B). We propose that there is an ancestral ontogenetic relationship between the inferred nervous system architecture observed in *C. capulus* and the LNN identified in *M. leidyi*. The occurrence of a comb nerve is more frequent than the LNN across these species (Fig. 4B), rejecting a possible retention of an LNN across stem and crown ctenophores. We propose below three possible and testable evolutionary scenarios that may explain the evolution of ctenophore neural architectures (Fig. 4C; see Discussion).

## Discussion

Understanding how the earliest animal nervous system architectures evolved remains a central challenge in biology, and ctenophores offer a uniquely informative perspective on this topic. In our present study, we mapped the expression patterns of the neuropeptide precursors *ML02212a* and *ML199816a* in *M. leidyi*. We revealed previously unrecognized anatomical and molecular complexity in the developing ctenophore nerve net, including potential functional specialization following segregation of the neuropeptide expression. We identified two principal features of comb-row neural organization, the ladder-like nerve net (LNN) and the comb nerve, that are recurrently reported in both fossil and extant ctenophores (15,25,33,39,40). These findings suggest that ctenophore neural circuits are more complex than currently appreciated.

Recent studies have demonstrated the preservation of nervous system patterns in Cambrian fossils (18–23), including ctenophores (25), where they are preserved as carbonaceous traces. Our re-examination of the carbonaceous traces of the Cambrian ctenophore *C. capulus* (ROMIP 514339) revealed a ladder-like arrangement at the comb area, which connects aborally with a ring-like structure. The transverse and longitudinal traces of the comb area resemble the LNN of one-week-old cydippids (Fig. 1, 4A). The ciliated groove traces are consistent with the ciliated groove nerve of large cydippids (Fig. 2), whereas the aboral ring-like structure, detectable in adult *M. leidyi*, coincides with the aboral carbon-rich ring of *C. capulus* (Fig. 2, 3, 4A). These congruencies thus support the hypothesis that the carbonaceous traces of *C. capulus* represent preserved neurons. Ladder-like carbonaceous traces are also possibly apparent in the closely related but diminutive and less well preserved *Ctenorhabdotus campanelliformis* (25) (Fig. S4). Supposed comb nerves are observed to converge aborally (originally interpreted as an artefact of compression of opposing sides of the specimen into a single layer) and may instead reflect the edges of the ladder-like nervous system flanking the combs (Fig. S4). The identification of organic molecules that constitute carbonaceous traces in fossils, where possible, may shed light on the molecular composition of these structures and facilitate their classification.

Extending our analysis to other fossil and extant ctenophores reveals additional anatomical features of the Cambrian nervous system that are congruent with those observed in extant species (Fig. 4B). The longitudinal dark compressions at the center of the combs in *G. hemispherica* and *F. vesanus* (Fig. 4B) are interpreted as longitudinal comb nerves and proposed to be homologous to the giant axon of *E. dunlapae* (Fig. 4B) (15,25,27,33) identification of a comb nerve in *M. leidyi* is a significant finding, as a comb nerve has so far only been described in *E. dunlapae*. Comparison of the comb nerves in *M. leidyi* and *E. dunlapae* will require further characterization of their development, ultrastructure, and molecular identity to assess potential homology.

Although multiple evolutionary scenarios are plausible, our analysis favours the hypothesis that both the LNN and the comb nerve formed components of the ancestral neural architecture, with subsequent reorganization during ctenophore evolution and development (Fig. 4C). Under this hypothesis, (Scenario 1, Fig. 4C) the coexistence of LNN-like architectures and discrete comb nerves in Cambrian fossils reflects retention of an ancestral condition rather than independent acquisition. This scenario interpretation is corroborated by our developmental data from *M. leidyi*, in which an initially widespread LNN undergoes topological rearrangement and condensation into the comb nerve. Such developmental transformations provide a plausible mechanism by which both structures could be inherited from a common ancestral condition while giving rise to the diversity of adult neural architectures observed among extant ctenophores. Alternative hypotheses remain viable but require additional assumptions. Interpreting the LNN as the sole ancestral neural topology (Scenario 2, Fig. 4C) would necessitate multiple independent losses or secondary acquisitions of the comb nerve (Fig. 4C), whereas favouring the comb nerve alone as ancestral (Scenario 3, Fig. 4C) requires the repeated evolution of ladder-like architectures or having systematically misinterpreted their presence in the fossil record and is inconsistent with the developmental sequence observed in *M. leidyi*. Although preservational bias may account for the absence of a comb nerve in some fossils, the repeated occurrence of LNN-like structures across disparate fossil and extant lineages and body plans is less readily explained under these two latter scenarios. Our preferred hypothesis - that the LNN and the comb nerve are developmentally linked and can be expressed to different degrees in adult ctenophores - (Scenario 1, Fig. 4C) - is explicitly testable and subject to revision. Comparative analyses of neuropeptide expression patterns during development in *E. dunlapae* and other ctenophore species will be essential for evaluating the homology and developmental relationships of the LNN and comb nerves present in different modes. The future discovery of fossil taxa preserving both structures in clear association, or demonstrating transitional configurations, including developmental sequences, would further refine or potentially overturn our model.

By exploiting the exceptional preservation of Cambrian ctenophores, our study integrates fossil and neuroanatomical evidence to propose a parsimonious and testable framework for the early evolution of neural architectures. This perspective helps reconcile disparate fossil morphologies with developmental data from living taxa and provides new insights into the origins and diversification of animal nervous systems. Moreover, our findings suggest that neuronal condensation around locomotory structures reflects shared constraints on nervous system architecture. In organisms with repeated locomotory units, longitudinal tracts interconnected by transverse connections may provide an efficient solution for coordinating distributed motor activity while minimizing wiring length and maintaining rapid signal transmission. Together, these results indicate that such neural layouts represent recurrent architectural solutions imposed by body organization and coordination demands, rather than reflecting shared ancestry across animal lineages.

## METHODS

### Animal husbandry

Adults of *M. leidyi* were maintained in 25L Kreisels as described previously (44–46). Upon gamete collection, fertilized eggs were allowed to develop in 1L glass beakers with aeration for 5 to 7 days and fed with *Brachionus*, prior to fixation. To allow further development, 1-week-old cydippids were transferred to 10L Kreisels and fed daily with increasing amounts of *Brachionus*. Larger cydippids (approximately 0.5 cm) and young adults (approximately 1 cm) were maintained in 25L Kreisels and fed with *Artemia* and *Brachionus*.

### Fixation of specimens

Young cydippids (1-week-old refers to 7-to-10-day-old cydippids and 2-week-old refers to 14-to-15-day-old cydippids) were fixed as previously described (44). In brief, younger cydippids were collected into 5ml glass vials containing 2.5ml of artificial seawater (ASW, 27ppt, pH=8.2) and relaxed by rapidly adding 500μL of 1M MgCl_2_. Fixation was carried out by firmly adding 16% ice-cold RainX® (47) in ASW and incubating for 20 min on ice under constant rotation. Cydippids were postfixed in 3.7% ice-cold Formaldehyde solution in ASW for 20min on ice under constant gentle rotation. Fixed specimens were then washed in 1X Phosphate Saline Buffer (PBS, 137 mM NaCl, 2.68 mM KCl, 10.14 mM Na_2_HPO_4_, 1.76 mM KH_2_PO_4_, pH=7.4) with 0.1% Tween 20 (Sigma Aldrich) 3 times for 10 min, dehydrated stepwise in methanol, and stored at -20°C in 100% methanol.

Larger cydippids (approximately 0.5cm large) and young adults (approximately 1-to-1.5 cm) were fixed as the younger stages with the following modifications. Animals were collected in glass vials containing 2.5 ml of artificial seawater (ASW, 27ppt, pH=8.2) and relaxed by adding 500 μL of 1M MgCl_2_, dropwise. Fixation was conducted by adding ice-cold RainX® dropwise, for a final concentration of 37.5% in ASW. Vials were gently mixed by inversion and incubated on ice for 30 minutes under constant gentle rotation. Specimens were post-fixed in 4% ice-cold Paraformaldehyde solution in 1X PBS for 1 hour on ice, under constant gentle rotation. Following fixation, specimens were washed in 1X PBS with 0.1% Tween 20 (Sigma Aldrich), 3-4 times quickly, then 3 times for 10 minutes, dehydrated stepwise in methanol, and stored at - 20°C in 100% methanol.

### Whole-mount *in situ* hybridization chain reaction and immunostaining

I*n situ* HCR v3.0 with split initiator probes was performed using probe sets (ML02212a, ML199816a), amplifiers, and buffers obtained from Molecular Instruments (https://www.molecularinstruments.com) (48). Samples were rehydrated in 1X PBS-0.1%Tween 20, and HCR was performed as previously described for smaller specimens (44) with the following modifications. Following rehydration, larger specimens (large cydippids and adults) were transferred to 2 ml tubes in approximately 500 μL of PBS-0.1% Tween 20. Larger samples were pre-washed with 250 μL of probe hybridization buffer (Molecular Instruments) and then incubated with fresh probe hybridization buffer for 1 hour at 37° C. Probes were used at a concentration of 16nM for all samples, accounting for the larger volumes of the larger samples, and incubated overnight for smaller specimens, and for approximately 48 hours for larger specimens at 37°C. Following probe incubation, samples were washed in probe wash buffer 4 times for 15 minutes at 37°C, then 5 times in 5X saline sodium citrate buffer (RNase-free 20X SSC, Invitrogen CAT AM9765) containing 0.1% Tween 20 (5XSSCT). Samples were pre-amplified in amplification buffer (Molecular Instruments) for 30 minutes. Hairpins were heated at 95°C for 90 seconds and allowed to cool down for 30 minutes at room temperature in the dark. Amplification buffer containing 60 nM hairpins, and the monoclonal anti-α-tubulin antibody (T6074, SIGMA Aldrich) at a concentration of 1:100 was added, and samples were incubated overnight in the dark at room temperature. Following amplification, samples were washed 3 times for 5 minutes in 5X SSCT, followed by an additional wash for 30 minutes and one for 5 minutes in 5X SSCT. Nuclear staining (Hoechst 33342 10mg/ml ThermoFischer Scientific, diluted 1:2000) and secondary antibody (Alexa-488 conjugated goat anti-mouse IgG, ab150117 Abcam, diluted 1:200) solutions were prepared in 5X SSCT, added to the samples, and incubated overnight at 4°C in the dark. Finally, samples were washed 3 times in 5X SSCT for 10 minutes and 3 times in 5X SSC for 10 minutes and mounted in SlowFade Glass ™ soft-set antifade mounting medium. The protocol combining HCR and immunostaining was adapted from (49).

### Confocal microscopy

Images of fixed samples were acquired with an Olympus FV3000 confocal laser scanning microscope; the Z-stack step size was set to 0.20 μm to image the nerve net. Maximum-intensity projections were obtained using Fiji (50), and contrast and brightness were adjusted to enhance the visualization of gene expression.

### Image analysis of *Ctenorhabdotus capulus*

Photographs of the *C. capulus* fossil were taken by Jean-Bernard Caron at the Royal Ontario Museum using cross-polarized light, backscatter electron (BSE) imaging, and scanning electron microscopy. Images of the whole specimen and higher magnification of specific areas were analyzed, and interpretive drawings were obtained in Fiji (50). The BSE images focusing on the aboral end and comb areas were thresholded for pixel intensity. Binary maps were obtained by weighting the dark pixels against the background. The maps were further interpreted by superimposing digital masks onto the pixel-dense regions estimated in the previous step. We obtained an interpretative drawing highlighting the darkened areas associated with the aboral region and comb rows. The backscatter electron images (BSE) focusing on the aboral end and comb areas (Fig. 4D, E) were thresholded based on pixel intensities against the background. This yielded binary maps of carbon-rich deposits (Fig. 4D’, E’), which were further interpreted by superimposing digital masks onto the pixel-dense regions (Fig. 4D’’, E’’).

## ACKNOWLEDGEMENTS

We thank Jean-Bernard Caron (Royal Ontario Museum) for generously sharing digital images of *Ctenorhabdotus capulus*. We thank Burkhardt lab members Ruth Styfhals and Brian Wehner for providing feedback on the manuscript and Eva-Lena Nordmann for assistance with the preparation of the supplementary video. This work was supported by the Michael Sars Centre core budget and the European Research Council Consolidator Grant (101044989, “ORIGINEURO”).

