## Supplementary Information for "Ancient nervous system architecture in a living ctenophore"

### SUPPLEMENTARY MATERIALS
### FOR

### Ancient nervous system architecture in a living ctenophore

Anna Ferraioli[1], Paula Miramón-Puértolas[1], Paula Altenkirch[1], Alexandre Jan[1],
Jeffrey Colgren[1], Jakob Vinther[2*], Pawel Burkhardt[1*]

[1]Michael Sars Centre, University of Bergen, Norway
[2]University of Bristol, United Kingdom

*Corresponding authors

**This file includes:**
Figs. S1 to S4
Movie S1

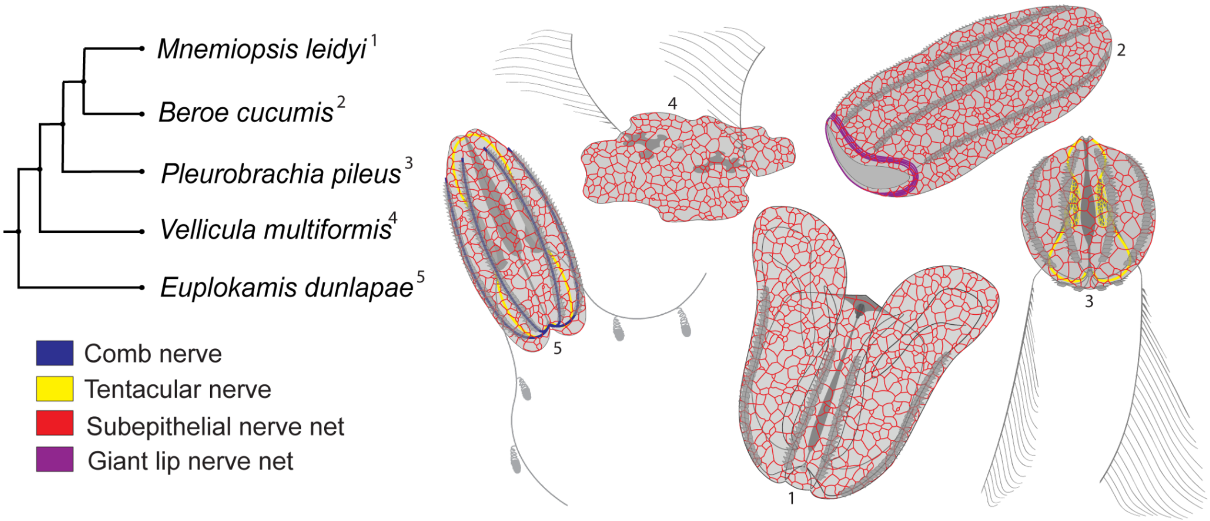


**Figure S1. Diversity of ctenophore nervous systems.** Phylogenetic relationships among representative ctenophore species with corresponding diagrams highlighting key neural architectures. While all ctenophores possess a polygonal subepithelial nerve net, *Euplokamis dunlapae* uniquely exhibits a comb nerve running beneath the comb plates. Both *E. dunlapae* and *Pleurobrachia pileus* possess a tentacular nerve. In *Beroe cucumis,* a prominent giant lip nerve encircles the oral region.

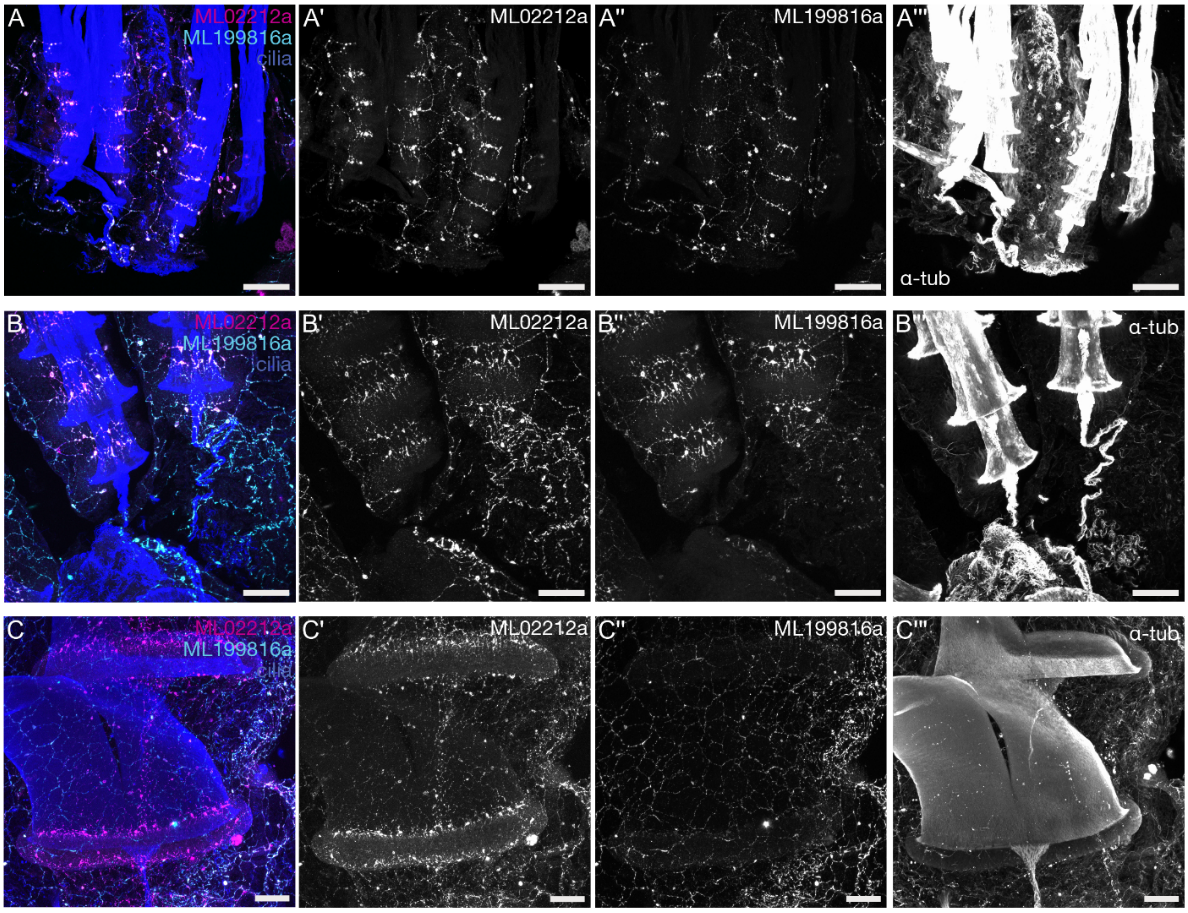


**Figure S2. Association of comb cilia and nerve net in *Mnemiopsis leidyi.*** HCR of *ML02212a* and *ML199816a* combined with tubulin staining (cilia). (A) Approximately one-week-old, lateral view of comb rows. (B) Two-week-old cydippid, top view of the comb rows, including the aboral organ and the ciliated grooves. (C) Transitioning adults (tentacles have not yet detached, but lobes are fully formed), lateral view of two comb plates, interconnected by the interplate ciliated nerve. Scalebars are 50 μm.

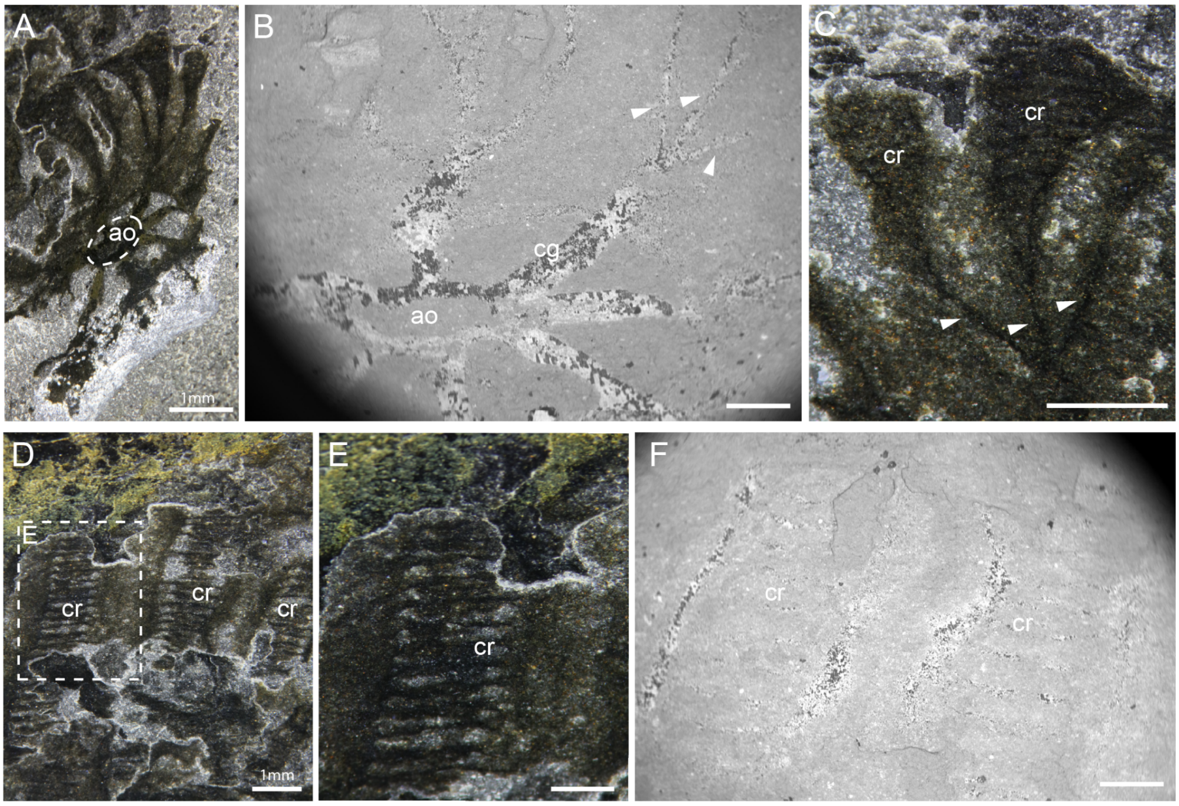


**Figure S3.** ***Ctenorhabdotus capulus*** **ROMIP 51439 morphology.** (A) *C. capulus* aboral organ region and part of the comb row photographed using cross-polarized light, showing the aboral organ fold. (B) Scanning electron microscopy backscatter of the aboral organ region and the narrow strands (white arrowheads) connecting with the comb rows. (C) Narrow strands (white arrowheads) and proximal comb row area (cr) photographed with cross-polarized light. (D) Comb row area photographed with cross-polarized light. (E) Higher magnification of (D). (F) Scanning electron microscopy backscatter of the comb row area. ao: aboral organ; cg: ciliated groove; cr: comb rows. Scalebars are 500 μm unless otherwise specified. Original fossil images are courtesy of J. B. Caron, Royal Ontario Museum.

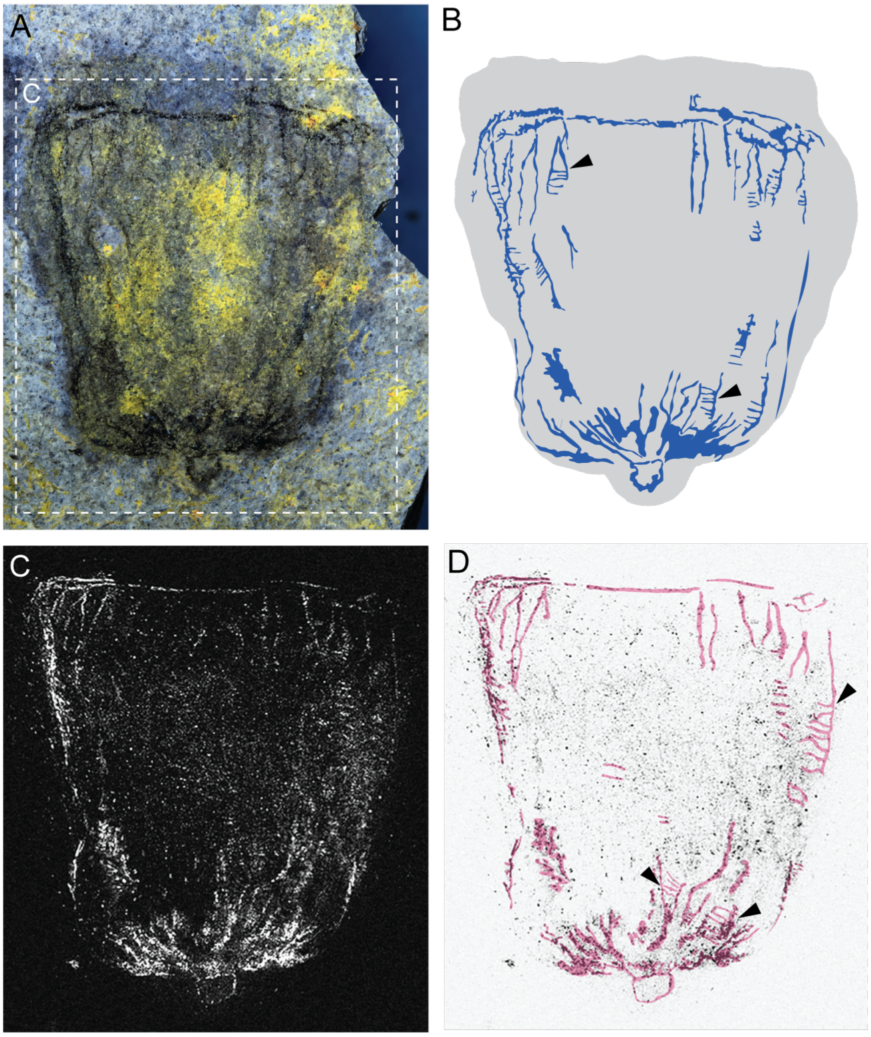


**Figure S4. Potential ladder-like patterns in *Ctenorhabdotus campanelliformis.*** (A) Cross-polarized light image of *C. campanelliformis* from (25). (B) Interpretative drawing based on the carbon-rich deposits in (A). (C) BSE imaging of (A) as in (25). (D) Interpretative drawing based on the carbon-rich deposits in (C). Ladder-like patterns can be found in certain areas in the oral and aboral areas (black arrowheads). Oral triangle-like patterns may therefore correspond to nerves framing the oral end of the combs.
